# Bridging genomic gaps: A versatile SARS-CoV-2 benchmark dataset for adaptive laboratory workflows

**DOI:** 10.1101/2024.04.24.587375

**Authors:** Sara E. Zufan, Louise M. Judd, Calum J. Walsh, Michelle L. Sait, Susan A. Ballard, Jason C. Kwong, Timothy P. Stinear, Torsten Seemann, Benjamin P. Howden

## Abstract

Genomic sequencing’s adoption in public health laboratories (PHLs) for pathogen surveillance is innovative yet challenging, particularly in the realm of bioinformatics. Low- and middle-income countries (LMICs) face increased difficulties due to supply chain volatility, workforce training, and unreliable infrastructure such as electricity and internet services. These challenges also extend to high-income countries (HICs) where bioinformatics is nascent in PHLs and hampered by a lack of specialized skills and computational infrastructure. This underlines the urgency for flexible and resource-aware strategies in genomic sequencing to improve global pathogen surveillance. In response to these challenges, the present research was conducted to identify and analyse key variables influencing the quality and accuracy of amplicon sequence data. An extensive benchmark dataset was developed that encompassed a diverse collection of isolates, viral loads, primer schemes, library preparation methods, sequencing technologies, and basecalling models, totalling 750 sequences. This dataset was analysed with bioinformatic workflows selected for varying levels of technical capacity. The evaluation focused on quality metrics, consensus accuracy, and common genomic epidemiological indicators. The analysis uncovers complex interactions between multiple parameters in laboratory and bioinformatic processes. emphasising resource-constrained PHLs, practical guidelines are proposed. Insights from the benchmark dataset aim to guide the establishment of specific laboratory and bioinformatics protocols for amplicon sequencing in these settings. The findings can also be used to guide the creation of specialised training curricula, further advancing genomic equity. The benchmark dataset itself allows laboratories to customise and evaluate workflows, catering to their distinct requirements and capacities. Such a holistic approach is imperative to build the capacity to monitor pathogens worldwide.

**Author summary:** This study marks a step toward equity in the field of pathogen genomics, especially for resource-constrained PHLs. It develops and evaluates a comprehensive amplicon sequencing benchmark dataset, offering vital insights for PHLs engaged in genomic surveillance. In particular, the study finds that the choice of basecaller model has a minimal impact on the quality and accuracy of consensus sequences derived from ONT data, which is crucial for labs with limited computational resources. It also highlights the effectiveness of longer amplicons in ensuring consistent coverage and reducing amplicon dropouts at higher viral loads. While Illumina remains a gold standard for data quality, the combination of the Midnight primer scheme with ONT’s Rapid library preparation is shown to be a viable alternative, reducing costs, procedural complexity, and hands-on time. The study synthesises these findings into practical guidelines to aid in the development of amplicon sequencing workflows for SARS-CoV-2 with implications for other pathogens.

## Introduction

The COVID-19 pandemic has demonstrated the transformative role of genomics in public health, transitioning from a predominantly academic tool to an essential component in pathogen surveillance and public health strategy. The integration of genomic data into public health frameworks has profoundly influenced global and regional surveillance of SARS-CoV-2 [1, 2], improving our understanding of the dynamics of virus transmission [3], evolutionary patterns [4], and the emergence of novel variants [5]. This information has been instrumental in shaping informed and effective public health responses [6].

The World Health Organization’s (WHO) call for widespread genomic sequencing to strengthen disease surveillance highlights its critical role in global health [7, 8]. However, there is a stark contrast in the sequencing capacity between high-income countries (HICs), which report more sequences per capita, and low- and middle-income countries (LMICs), despite the commendable efforts in some African nations [9–11]. Moreover, the urgent demand for genomic sequencing in public health from the pandemic has exposed several challenges in all economic contexts. In LMICs, limited access to genomic technologies, lab infrastructure, supply chain reliability, training of the workforce, and data quality assurance prevail [12, 13]. In contrast, HIC PHLs face the need to expand their bioinformatics personnel and improve their computational infrastructure to handle the surge in genomic data [14]. These opposing sets of challenges emphasise the need for versatile and resilient genomic surveillance strategies that are effective in varying economic and resource contexts.

In addressing these challenges, the potential of benchmark datasets in public health genomics, particularly for quality assurance, should be explored. These datasets, which offer high-quality genomic sequences and annotations, are crucial for the validation of bioinformatic tools and methods used in outbreak surveillance [15, 16]. By engaging with these datasets, laboratories can critically evaluate and adapt sequencing workflows to their resource availability, establishing nuanced quality control parameters. However, existing benchmark datasets for SARS-CoV-2 often cover only a limited range of variables (e.g. primer scheme, sequencing platform). For example, recent interlaboratory validation studies have highlighted discrepancies in variant calling at mixed allele sites, such as those arising from intrahost variation, attributed to platform-specific bioinformatic workflows [17, 18]. The multitude of variables inherent in laboratory and bioinformatic workflows underscores the need for more comprehensive datasets that can more effectively address the wide range of challenges encountered in diverse global contexts.

This study provides a benchmark dataset for SARS-CoV-2 amplicon sequencing, reflecting a wide range of isolates, primer schemes, sequencing technologies, and bioinformatic approaches. Our analysis determines how different laboratory and bioinformatic workflows affect data quality and consistency. The insights gained aim to guide the formulation of evidence-based practices for effective amplicon sequencing across all PHLs, regardless of resource availability. This work supports the advancement of training initiatives around the world, promoting equitable genomic technology proficiency for enhanced global health surveillance.

## Materials and methods

### Sample preparation

This benchmark dataset’s contrived specimens were sourced from five isolates, courtesy of the Victorian Infectious Disease Reference Laboratory (VIDRL) (S1 Table). Their isolation and preparation adhered to established protocols previously used for SARS-CoV-2 interlaboratory assessments [19, 20]. Briefly, the negative sample matrix was prepared by pooling 1 mL of viral transport medium (VTM) from 300 throat and deep nasal swabs, all previously confirmed negative by the Aptima^®^ SARS-CoV-2 assay. This pooled VTM was divided into 10 aliquots and retested to verify negativity. Subsequently, 3 ml of this confirmed negative matrix was mixed with 55 µL of heat-killed viral stock in a sterile 5 mL tube, followed by inverting five times. A 500 µL sample was then transferred to a Panther Fusion specimen lysis tube for analysis using the Hologic Panther Fusion SARS-CoV-2 PCR assay. To simulate varying viral concentrations, the samples were serially diluted 1:10 in triplicate using the remaining negative matrix and quantified with the SARS-CoV-2 PCR E-gene assay.

### Amplicon sequencing

Specimens were processed using multiplexed amplicon sequencing with the ARTIC V4 [21] and Midnight [22] primer schemes. Libraries for Illumina sequencing were prepared using the Nextera XT kit. For Nanopore sequencing, libraries were prepared using the ONT Native Barcoding Kit (EXP-NBD104) (hereafter referred to as Ligation) and the ONT Rapid Barcoding Kit (SQK-RBK004) (hereafter referred to as Rapid). Sequencing was performed on the Illumina iSeq and Nanopore GridION platforms, using MinION flow cells (S2 Table). A breakdown of the entire workflow is depicted in Fig. 1.

**Fig 1.**
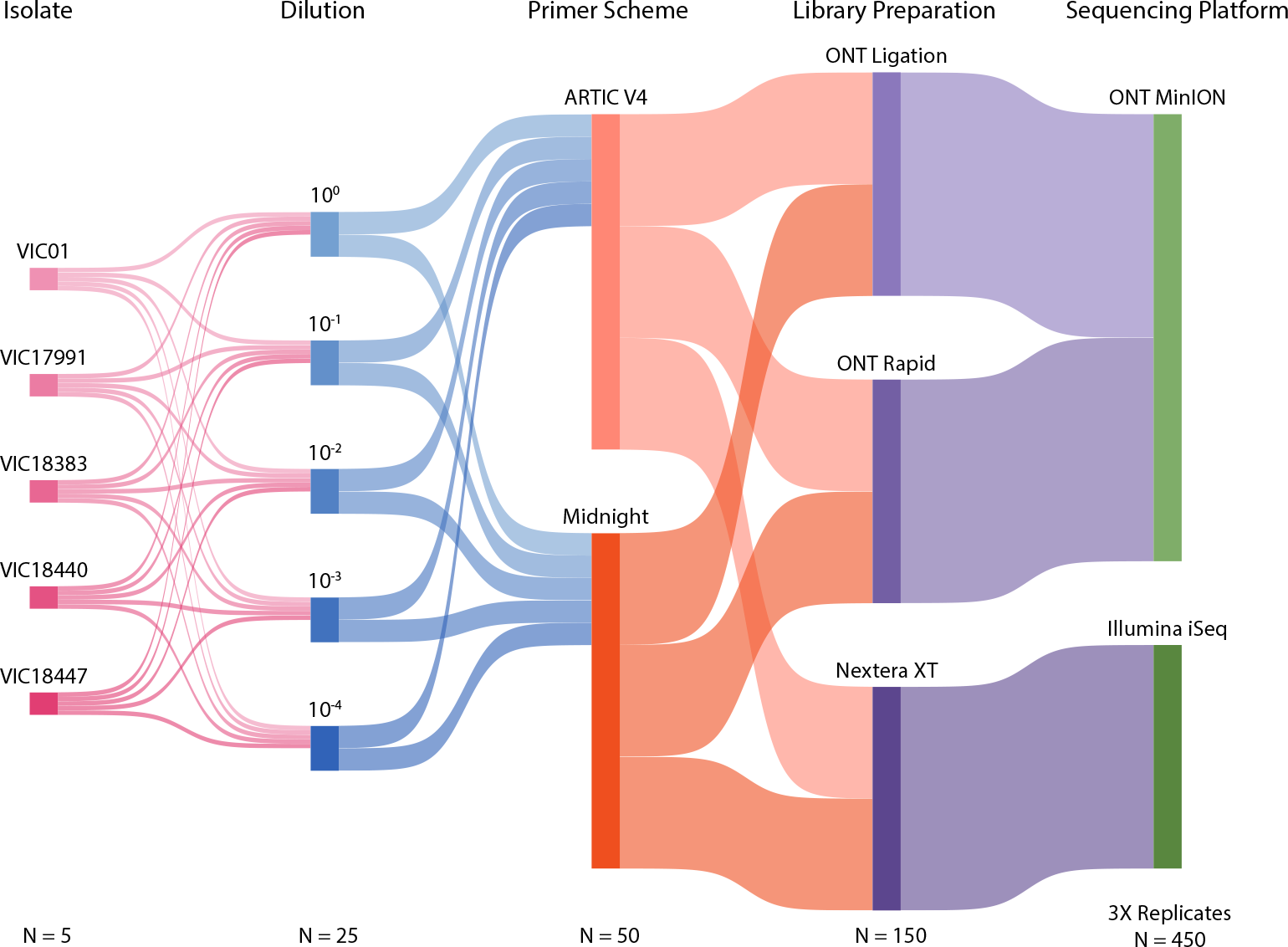
Sample preparation workflow. Sankey diagram depicting the sample preparation workflow for the SARS-CoV-2 amplicon sequencing benchmark dataset. Beginning with five distinct isolates, each undergoes serial dilution before proceeding through two primer schemes: ARTIC V4 and Midnight. Subsequent library preparation is conducted using ONT Ligation and ONT Rapid library preparation for sequencing on ONT MinION, and Nextera XT for the Illumina iSeq platform. Technical replicates were performed for all samples totaling 450 sequences. Additionally, positive and negative controls were processed concurrently.

### Bioinformatic workflows

Established methods commonly utilised in PHLs were employed for the generation of consensus sequences. Short-read data were processed using `iVar` software (version 1.4.2) [23], where primer trimming and consensus sequence generation were conducted. Parameters were set at a quality threshold of 20 (-q 20), a variant calling threshold of zero (-t 0), and a minimum depth of 10 (-m 10), conforming to standards widely accepted in the field [16]. This approach aligns with web-based pipelines [24] and the methods prevalent in PHLs [1], ensuring compatibility with current practices.

Nanopore reads were basecalled using `Guppy` (v6.2.11) super high accuracy (hereinafter referred to as ‘Super’) and fast (hereinafter referred to as Fast) models, representing variability in computational infrastructure. Basecalled, demultiplexed reads were processed using the `wf-artic` (v0.3.30) workflow [25]. This workflow, available through ONT’s EPI2ME bioinformatics platform, offers both a graphical user interface (GUI) and command-line functionality. `wf-artic` is a derivative of the widely utilised artic pipeline [26], with notable modifications. These include optimised primer trimming for fragmented reads, a common occurrence in transposase-based rapid barcoding systems (https://github.com/artic-network/fieldbioinformatics/issues/99).

### Performance evaluation

Quality metrics were collected using `ncov-tools` (v1.9.1) (https://github.com/jts/ncov-tools), QualiMap (v2.3) [27], and `nextclade` (v2.14.0) [28]. Metrics included read count, error rates, base quality, amplicon depth, breadth of coverage, lineage assignment, phylogenetic clustering, and consensus accuracy. Accuracy was evaluated by comparing the variants identified in specimen sequences with those in the primary isolate sequence (Table **??**). Within this framework, the variants detected in both the specimen and the primary isolate were classified as true positives (TP). Variants identified in the specimen but not in the primary isolate were considered false positives (FP). In contrast, variants identified in the primary isolate but absent in the specimen were deemed false negatives (FN). The following measures were used for statistical evaluation:

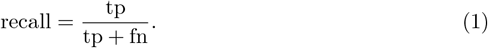

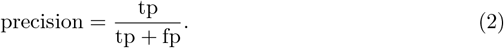

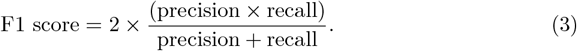

All subsequent statistical analyses were performed in R (v4.3.0) and `SciPy` (v1.11.4).

## Results and discussion

### Benchmark sample characteristics

The benchmark dataset was made up of contrived specimens *N* = 75, including two biological replicates in five lineages and dilutions, and one set of technical replicates performed in triplicate. TCID50 values ranged from 0.01 to 1000 (mean 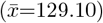; median 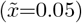; inter-quartile range (IQR)=9.97) (Fig. 2). Amplicon sequencing was performed using the ARTIC V4 and Midnight schemes, with library preparations from Nextera XT, ONT Ligation, and ONT Rapid kits, resulting in 450 samples for sequencing. Furthermore, ONT sequences were basecalled using Super and Fast models, totalling 750 sequences for comprehensive analysis.

**Fig 2.**
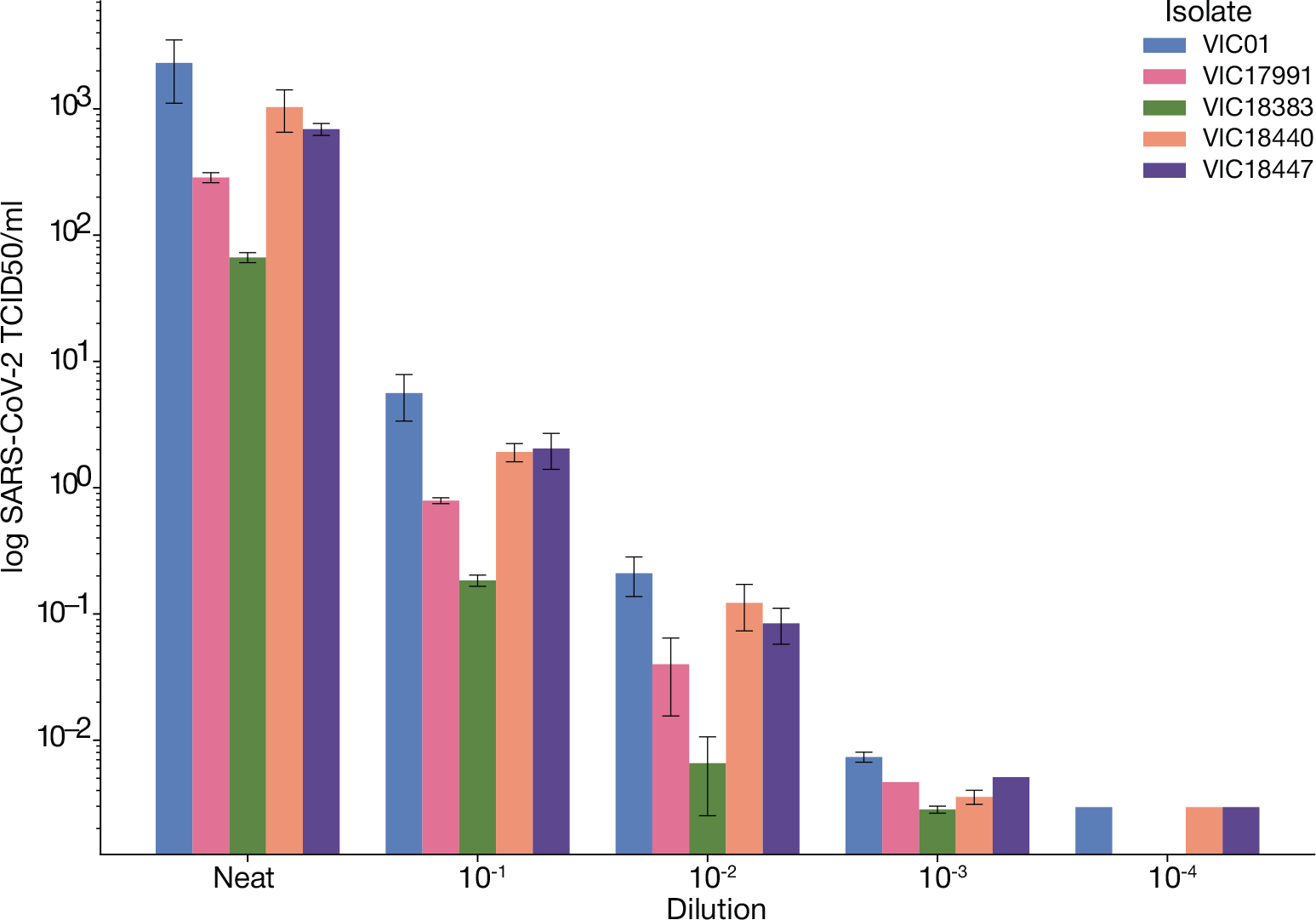
TCID50 estimates for contrived specimen. Quantitative assessment of SARS-CoV-2 viral titers across various dilutions and isolates prepared for this study. TCID50 values were extrapolated from Ct values obtained via the Panther Hologic fusion assay, guided by a predefined standard curve (S1 Fig.

### Basecaller performance

#### Nanopore run summaries

Each scheme and library preparation pair were run on an individual MinION flow cell. Initial pore scans ranged from 371 (18.12%) to 1714 (83.70%) active pores (Fig. 3A). Among the Midnight scheme runs, Ligation achieved a basecall pass rate of 74.22% with a total output of 65.92 Mb (Fig. 3B). In contrast, Rapid obtained a pass rate of 64.00% and a total output of 886.17 Mb, reflecting the low initial pore activity (Fig. 3). With respect to ARTIC V4, Ligation showed a notable basecall pass rate of 85.87% with a total output of 712.11 Mb. Rapid had the lowest pass rate of 28.02%. Considering the high initial pore activity, the low pass rate may be indicative of poor library purity.

**Fig 3.**
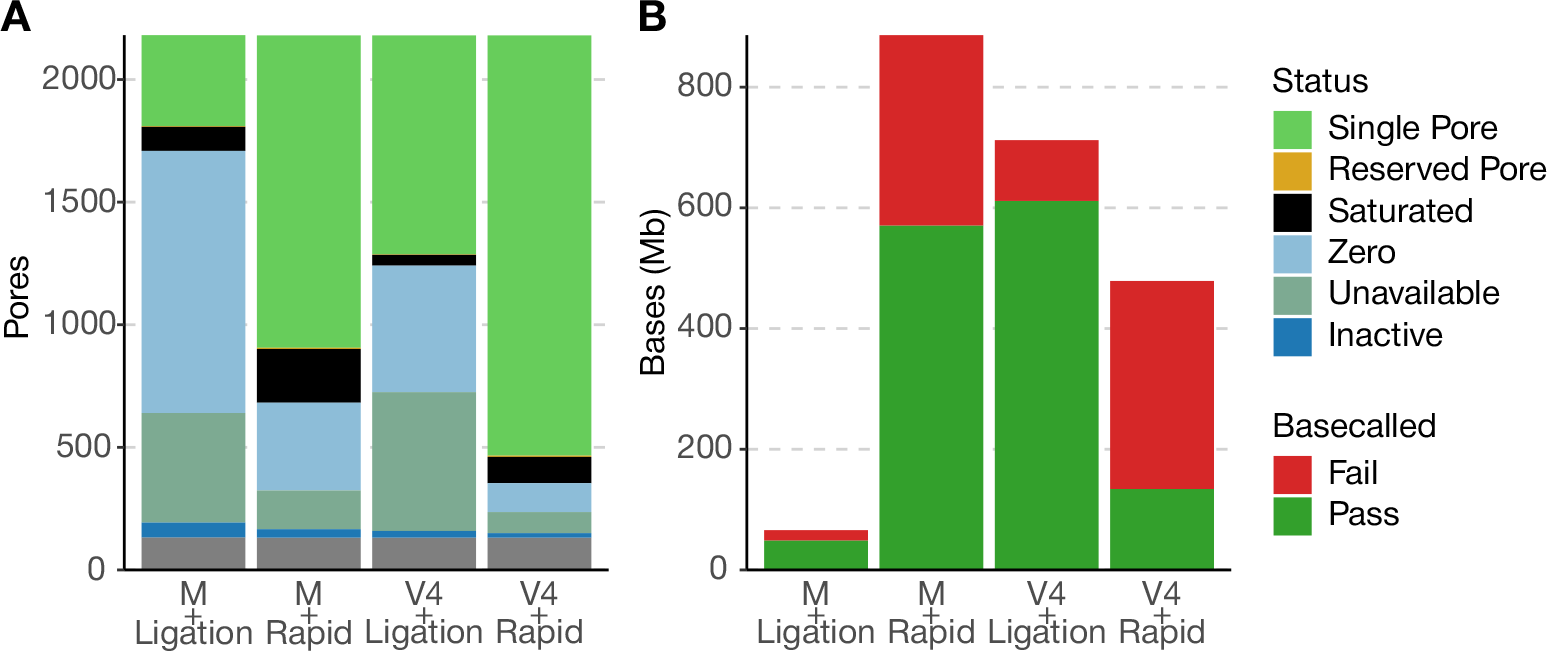
Comparative characteristics of Nanopore sequencing runs. A) Distribution of initial pore statuses across multiple runs, reflecting the operational efficiency of the flowcells used. B) Aggregate basecalling output for each run, categorized into bases that passed or failed the quality threshold (Q ≥ 7).

#### Basecaller performance

The ONT base caller models were evaluated for average base quality (avgBQ), mismatch count, and consensus accuracy. The Fast model showed a mean avgBQ of 25.09 with a median of 27.90 and an IQR of 10.90 (S2 FigC), while Super had a mean of 24.75, a median of 24.45, and a smaller IQR of 8.00, indicating a more consistent avgBQ. No significant difference in avgBQ was observed between Super and Fast (Wilcoxon rank sum, p = 9.654e-01). Stratification by library preparation (S2 FigB) revealed Rapid preparation improved avgBQ with Super 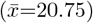 over Fast (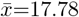; Wilcoxon rank-sum, p=3.313e-34), unlike Ligation. Primer scheme analysis (S2 FigA) showed that with Midnight, Super’s avgBQ 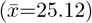 exceeded Fast’s (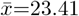; Wilcoxon rank-sum, p=1.160e-04), whereas with ARTIC V4, Fast outperformed Super (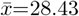 vs. 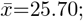; Wilcoxon rank-sum, p=6.709e-10). In particular, Midnight with Rapid was the only significant differential, favouring Super (Wilcoxon rank sum, p = 4.308e-26), aligning with the trend that Rapid avgBQ improves when using the Super model. Sparse Midnight with Ligation data likely influenced these scheme-specific divergences.

Despite variations in avgBQ across multiple strata, a regression analysis found no significant correlations between avgBQ and mismatches (S2 FigD) or accuracy (S2 FigE) for both Fast and Super models. Pearson’s correlation coefficients (*r*) for Fast showed negligible relationships ((r = −0.03)) for mismatches and percent consensus accuracy ((r = −0.03)). Similarly, Super only exhibited weak correlations ((r = 0.21)) for mismatches and precision ((r = −0.08)).

Furthermore, no statistically significant differences were observed in the number of mismatches (Mann-Whitney U test, p = 0.899) or in the percentage of accuracy (Mann-Whitney U test, p=0.896) between the Super and Fast models. In particular, the Fast model, while maintaining comparable performance, offers computational efficiency, making it suitable for resource-limited environments. Subsequent analyses will primarily focus on the Fast model data, aligning with the study’s objectives unless otherwise specified.

#### Benchmark dataset characteristics

Taking into account the importance of achieving a breadth of coverage ≥90% as a key QC threshold for the subsequent genomic epidemiological analysis [16, 18], the study evaluated consensus sequences against this standard. A total of 30 sequences met this criteria (6.67%). The Midnight primer scheme paired with Nextera XT library preparation attained the greatest genome completeness rate of 19.74% (n=15), closely followed by Midnight paired with Rapid, with 17.33% (n=13) (Fig. 4A). ARTIC V4 paired with Nextera XT showed comparable results, while ARTIC V4 paired with Ligation lag significantly, with only 2. 67% (n = 2) achieving the required coverage. Both Midnight paired with Ligation and ARTIC V4 paired with Rapid did not meet the ≥ 90% coverage threshold. Consequently, further analyses concentrated on the 10^0^ dilution series, as only one sample met the breadth threshold in subsequent dilutions (S3 Table).

**Fig 4.**
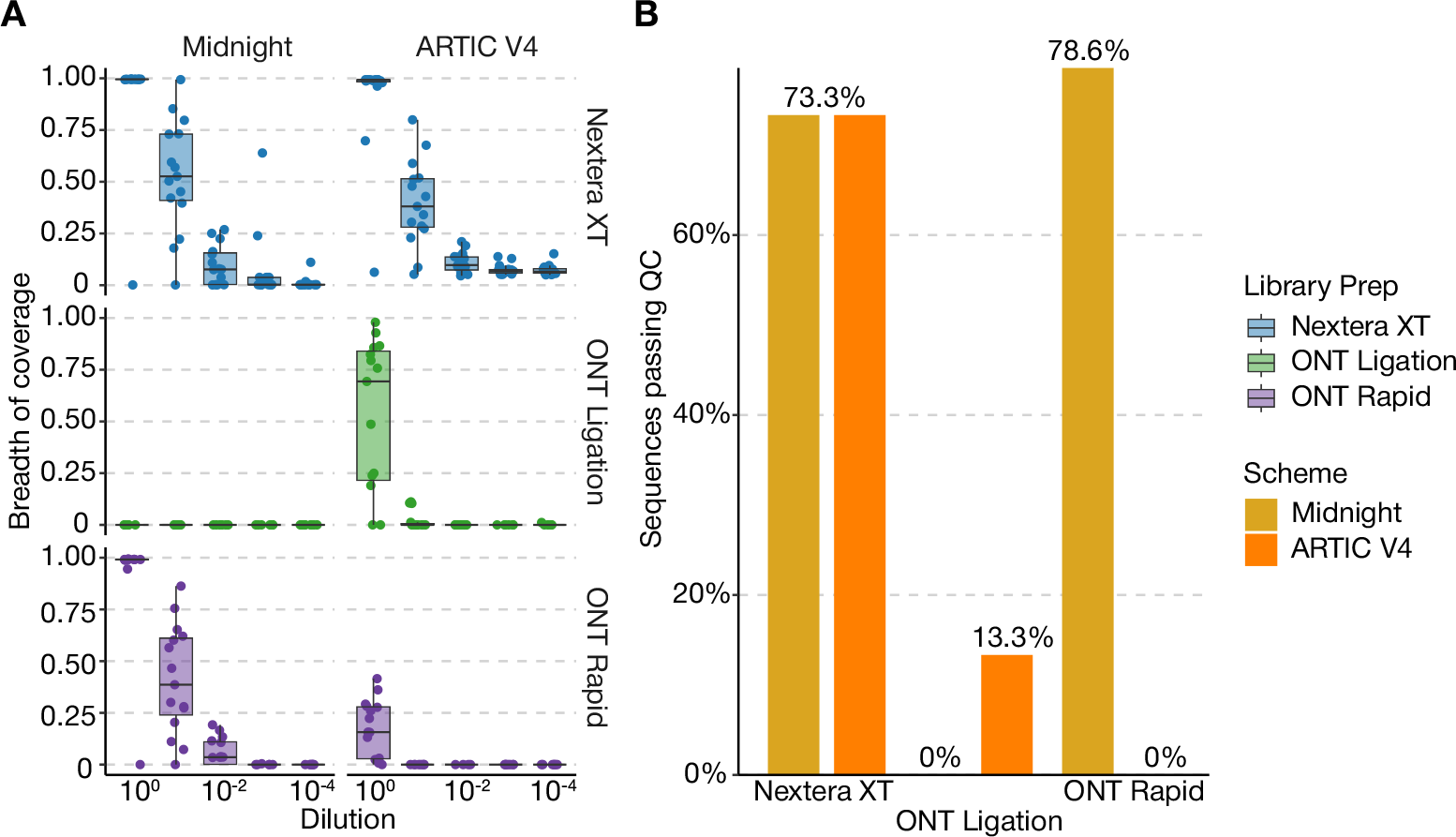
Characteristics of consensus sequences. Excluding samples processed with the the Super-basecalling model, which showed negligible influence on quality control (QC) metrics, panel A illustrates the breadth of coverage across serial dilutions. Here, individual data points denote the breadth of coverage for each sample, with boxplots providing a summary of the overall data distribution. Panel B examines the proportion of sequences that met the `nextclade` QC criteria, segregated by library preparation method and primer scheme, specifically focusing on the 10^0^ dilution subset.

In this narrowed scope of the 10^0^ dilution series, the Pass rate in `nextclade` QC status became a focal point. Here, Pass was defined as receiving an overall QC status of “good” or “mediocre”. Both Midnight paired with Nextera XT and ARTIC V4 paired with Nextera XT demonstrated a Pass rate of 73.33%, indicating their efficiency. In contrast, Midnight paired with Rapid outperformed slightly with a Pass rate of 78.57%, whereas ARTIC V4 paired with Ligation exhibited a significantly lower rate of 13.33% (Fig. 4B).

### Benchmark dataset performance

#### Breadth of coverage

The performance of various sequencing schemes and library preparation methods was evaluated, with a focus on the breadth of coverage at different dilution levels. The analyses were concentrated in dilutions 10^0^ and 10^1^ due to the low viral concentrations in the prepared sample.

In the 10^1^ dilution analysis, the Midnight scheme paired with Nextera XT showed a high mean coverage of 76.06% (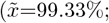; IQR=48.02%), while Midnight with Rapid had a 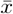 coverage of 68.87% (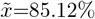; IQR=65.14%). ARTIC V4 paired with Nextera XT reported a 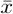 coverage of 65.38% (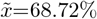; IQR=62.74%), and with Ligation, it had the lowest 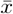 coverage of 47.27% (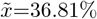; IQR=71.04%).

At the 10^0^ dilution, the results differed significantly. Midnight with Nextera XT showed a high and consistent coverage, with a 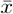 of 99.51% (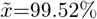; IQR=0.04%). Similarly, Midnight with Rapid also demonstrated high coverage (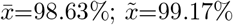 IQR=0.18%). In ARTIC V4 sequencing, Ligation exhibited a 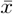 coverage of 62.85% (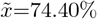, IQR=55.46%), while Nextera XT had a mean coverage of 88.69% (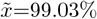, IQR=1.23%).

The stratified analysis revealed significant variations in the breadth of coverage ≥90% between different primer schemes and library preparation methods. For example, Midnight with Nextera XT achieved 14. 67% high-coverage samples (n = 11/75) in the Nextera XT and Rapid preparations, compared to only 2.67% (n=2/75) high-coverage samples for ARTIC V4 with Ligation.

In general, these findings emphasise the significant impact of primer schemes and library preparation methods in achieving the high breadth of coverage needed for accurate genomic epidemiological analysis of SARS-CoV-2. The Midnight scheme consistently showed high coverage across different library preparations, while ARTIC V4 displayed more variability. This is in agreement with Freed *et al*., who found the length of the 1200 bp amplicon of Midnight to be optimal for uniform coverage across viral concentrations [22]. However, it should be noted that Midnight outperformed ARTIC V4 with higher viral loads (∼ TCID50 ≥ 5, or, *Cτ* ≤30) (S3 Fig). Furthermore, dilution 10^0^ generally led to higher and more consistent coverage, especially with Nextera XT, underscoring the importance of considering viral load when prioritising candidate samples for genomic sequencing.

#### Consensus sequence evaluation

Comparison of consensus sequence accuracy metrics across various primer schemes and library preparation methods indicated significant performance differences. The Midnight scheme, when used with Rapid library preparation, exhibited the highest metrics, demonstrating a recall of 59. 27%, precision of 92. 95%, and an F1 score of 65. 44%. In contrast, the combination of Midnight with Nextera XT displayed moderate performance, with a recall of 55. 36%, precision of 78. 48%, and an F1 score of 59. 03%.

The ARTIC V4 scheme showed more variability in its performance. Together with Ligation, it achieved a recall of 49. 83%, precision of 93. 33%, and an F1 score of 57. 57%. However, when ARTIC V4 was combined with Nextera XT, it resulted in the lowest performance metrics among the evaluated groups, with a recall of 29. 33%, precision of 43. 74%, and a F1 score of 31. 97%. This relatively lower performance of the ARTIC V4 scheme with Nextera XT, particularly in samples with higher Cvalues *τ*, aligns with the findings of Mboowa *et al*., who reported that the ARTIC SARS-CoV-2 sequencing protocol on the Illumina MiSeq platform was sensitive and accurate to C*τ* of 24 in Ugandan settings [29]. This context highlights the importance of selecting appropriate sequencing protocols based on the characteristics of the samples being analysed.

At the 10^0^ dilution level, the Midnight scheme with Nextera XT achieved the highest scores, achieving perfect recall, precision, and F1-Score. The Midnight scheme with Rapid also showed high efficiency, with recall at 98.91%, precision at 100%, and F1-Score at 99.44%. ARTIC V4 with Ligation showed moderate results (recall = 66. 00%, precision = 100%, F1-Score = 72. 75%), with slight improvement when paired with Nextera XT (recall=88.73%, precision=91.67%, F1-Score=90.02%).

Combined, these results indicate that the Midnight scheme yields the greatest coverage and accuracy overall. Illumina continues to produce more accurate results, as has been continuously reported [30, 31]. However, read accuracy continues to improve with advances in ONT technology and basecalling models [32, 33].

Examining lineage assignment and phylogenetic clustering in our study reinforced the importance of stringent QC thresholds in genomic epidemiology. Despite variations in sequencing accuracy between different methods, 100% concordance was observed for both lineage assignment and phylogenetic clustering in all combinations with a coverage breadth greater than 80% (Fig. 5). Similarly, concordance was observed when `nextclade` designated overall QC status as “good” or “mediocre”. All pairs showing a steep decline in concordance when overall QC status was designated as “bad” (S4 Table).

**Fig 5.**
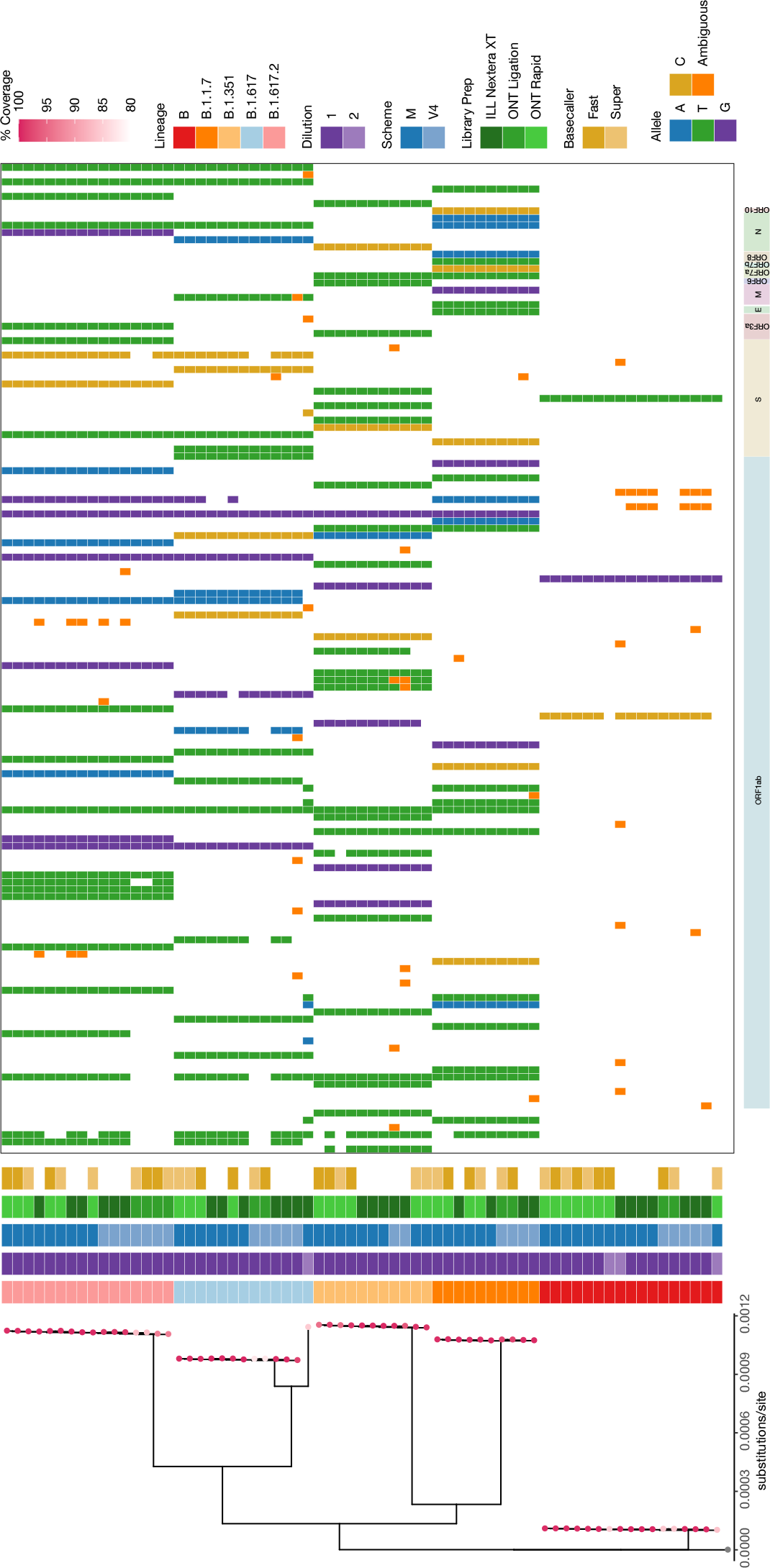
Clustering and characteristics of consensus sequences. Consensus sequences with breadth of coverage≥ 80% were selected for phylogenetic analysis in with `ncov-tools`. Tip colors denote breadth of coverage, followed by columns of sequence characteristics. The internal table show variants called colored by allele along the genome.

#### Limitations

Although it provides valuable information, this study is not without limitations. A primary challenge was the high C*τ* values of the specimens, indicative of low viral concentrations. This aspect posed difficulties across all sequencing methods, impacting their efficiency and consequently limiting the number of sequences available for thorough analysis.

The study’s challenges with two failed ONT runs, resulting in low yield and quality, limited the comprehensive evaluation of all primer scheme and library preparation combinations. This shortfall could bias interpretations due to missing data points. These issues, characteristic of ONT sequencing’s variable quality and yield [34, 35], are particularly pertinent for resource-constrained PHLs in developing effective sequencing workflows. Despite these constraints, the study’s findings are valuable, emphasising the need for adaptable sequencing methodologies to accommodate such inherent uncertainties in genomic sequencing.

This benchmark dataset, designed primarily for SARS-CoV-2, presents a flexible model suitable for various pathogens. The increasing use of multiplexed amplicon sequencing in pathogen surveillance, including Dengue [36], Mpox [37], and *Plasmodium falciparum* drug resistance and analysis of vaccine targets [38], underscores its importance. Our findings reveal a complex interplay of variables in the sequencing process, highlighting the need for meticulous quality assurance. Developing comprehensive benchmark datasets is crucial to ensure data quality in various settings, particularly in resource-constrained PHLs, and promotes genomic equity, thereby significantly improving global health surveillance.

### Conclusion

The comprehensive evaluation of primer schemes and library preparation methods in this study, using a robust benchmark dataset, offers essential guidance for resource-limited PHLs. Key findings include the minimal impact of basecaller model selection on the quality and accuracy of consensus sequences, which is advantageous for laboratories with limited computational resources. The research further reveals the benefits of longer amplicons at elevated viral concentrations, such as more uniform coverage and reduced susceptibility to dropouts, thus decreasing the need for frequent monitoring and replacement. Although Illumina remains the gold standard for data quality, the combination of Midnight primer schemes with ONT Rapid library preparation achieved comparable metrics. A synthesis of the findings, along with the existing literature, is presented in Table 1. This collated guide is designed to support the development and evaluation of SARS-CoV-2 amplicon sequencing workflows and can be adopted for other pathogens. In general, this study significantly contributes to the advancement of global genomic surveillance equity and the strengthening of public health strategies in the rapidly evolving domain of genomic sequencing.

**Table 1.**
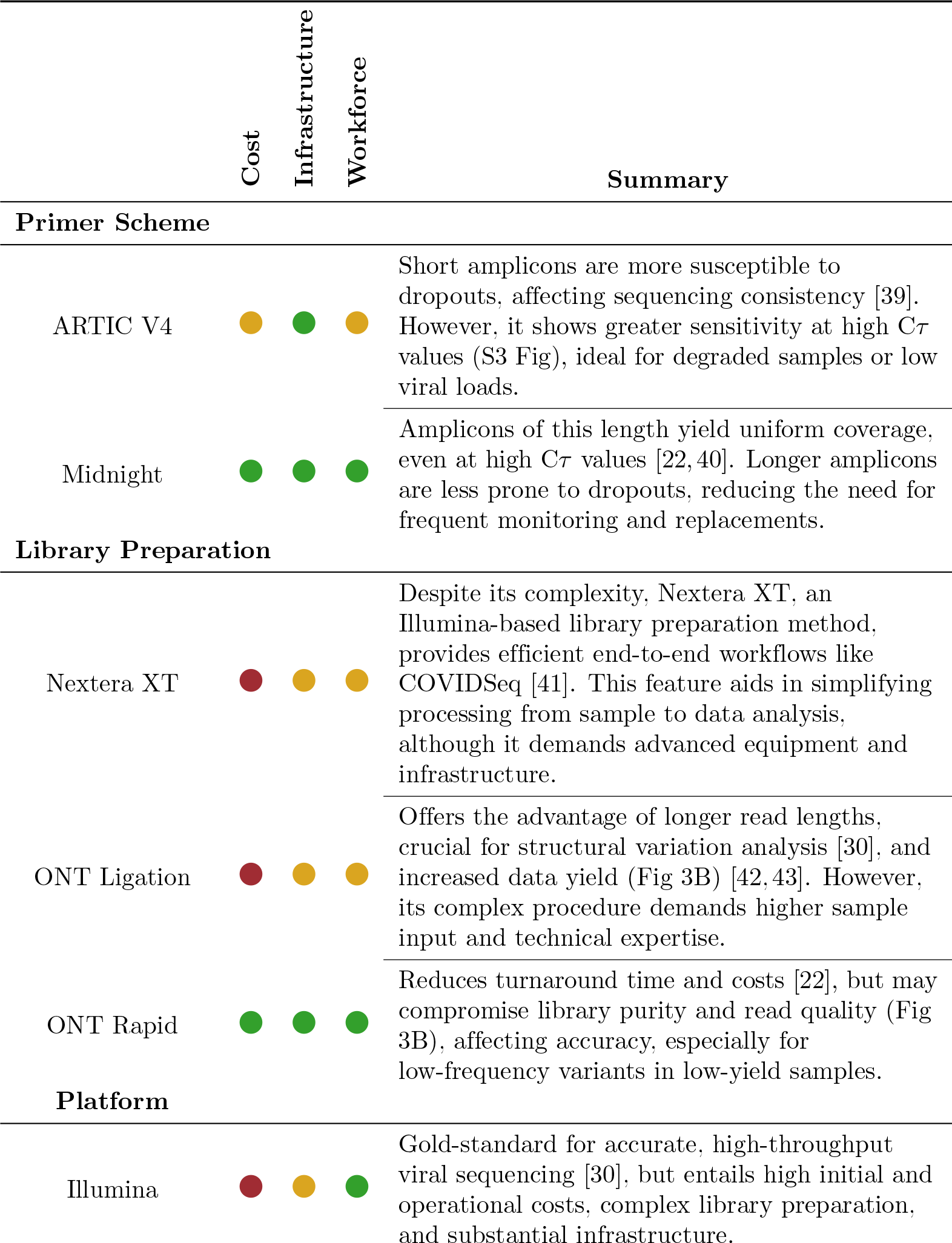
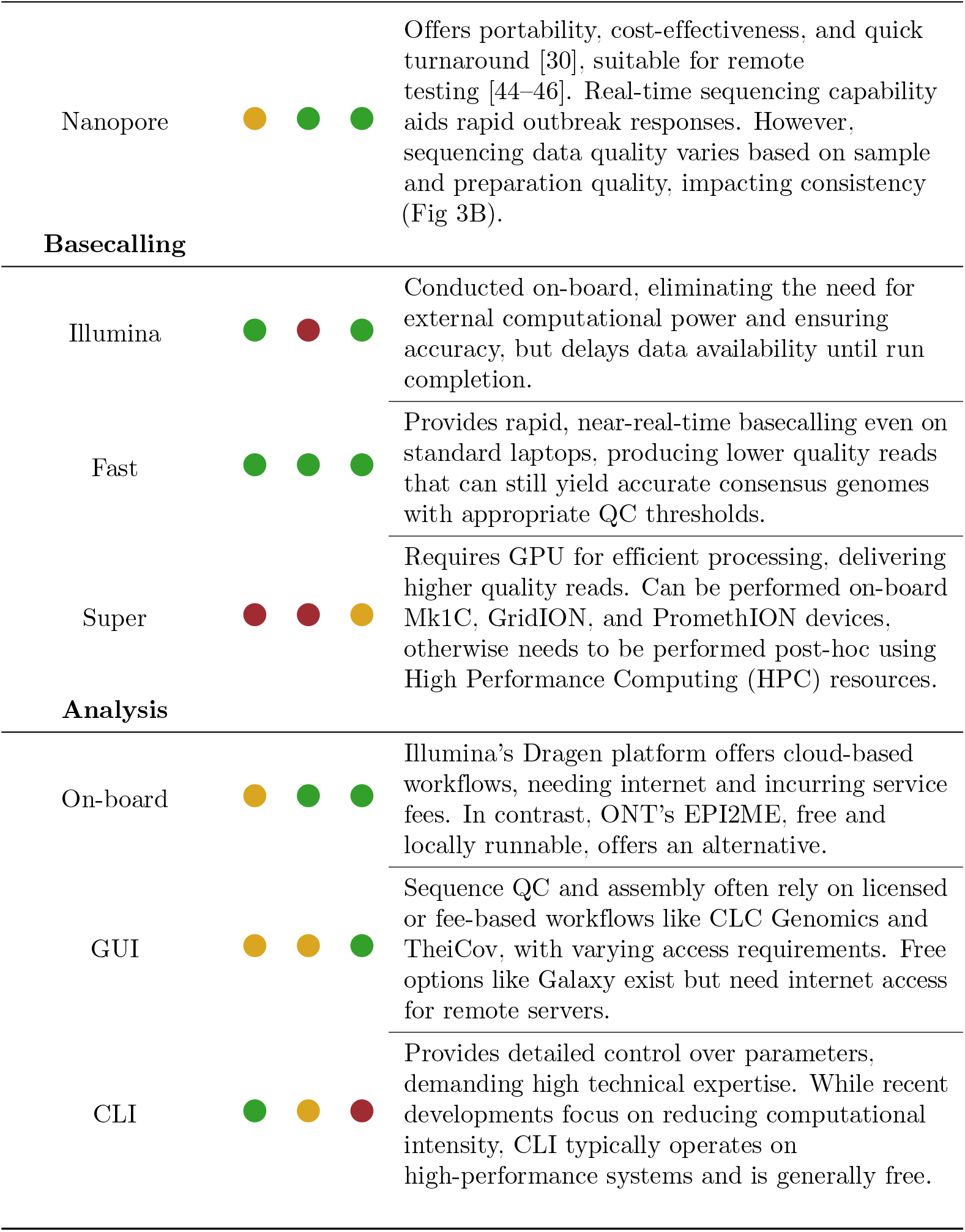
Guidelines for developing amplicon sequencing workflows in resource-constrained settings. The resource intensity is denoted as high 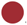, medium 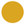, and low 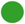.

## Acknowledgments

We acknowledge the support of the National Health and Medical Research Council, Australia, through their Partnership Grant (GNT1149991), and the Australian Government’s Medical Research Future Fund under the Genomics Health Futures Mission Flagships—Pathogen Genomics Grant (FSPGN000045) for this work. Additionally, S.E.Z. is a recipient of the Melbourne Research Scholarship from the University of Melbourne. Our gratitude extends to the Victorian Infectious Disease Reference Laboratory for supplying the viral isolates used in this study.

## Notes

### Competing Interest Statement

The authors have declared no competing interest.

