## Supplementary figures and images for "Bridging genomic gaps: A versatile SARS-CoV-2 benchmark dataset for adaptive laboratory workflows"

### Supplemental Figure 1

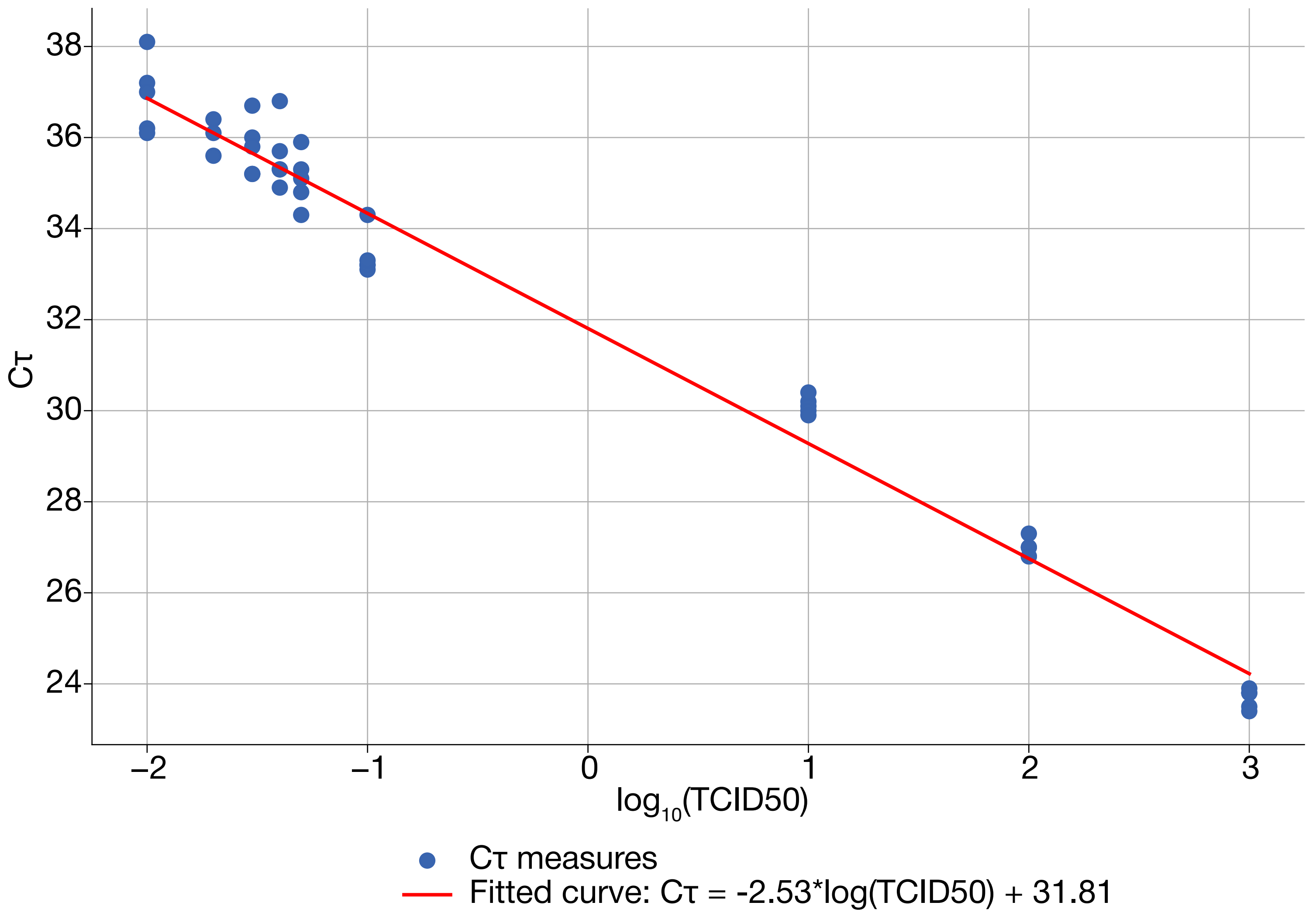

### Supplemental Figure 2

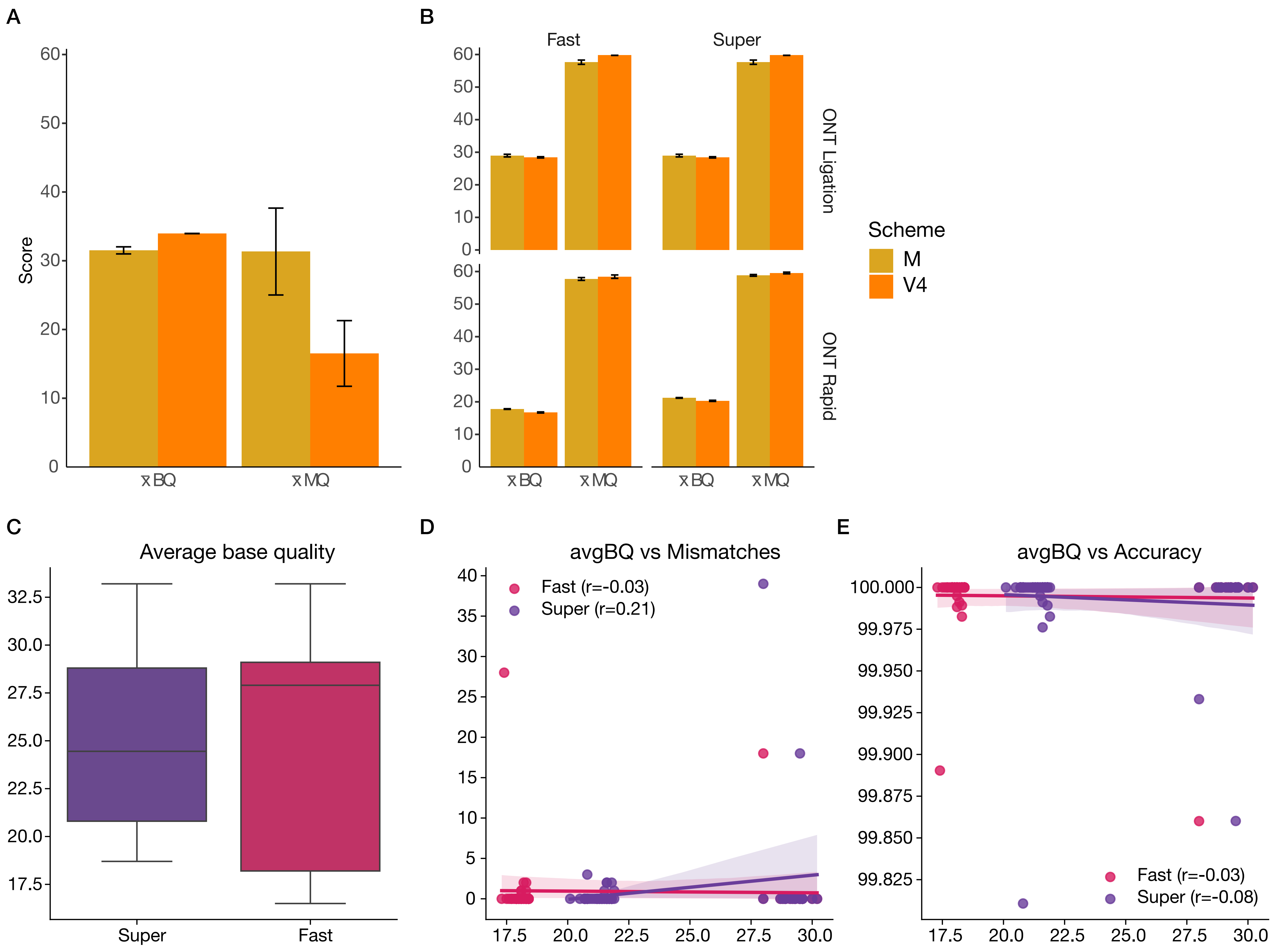

### Supplemental Figure 3

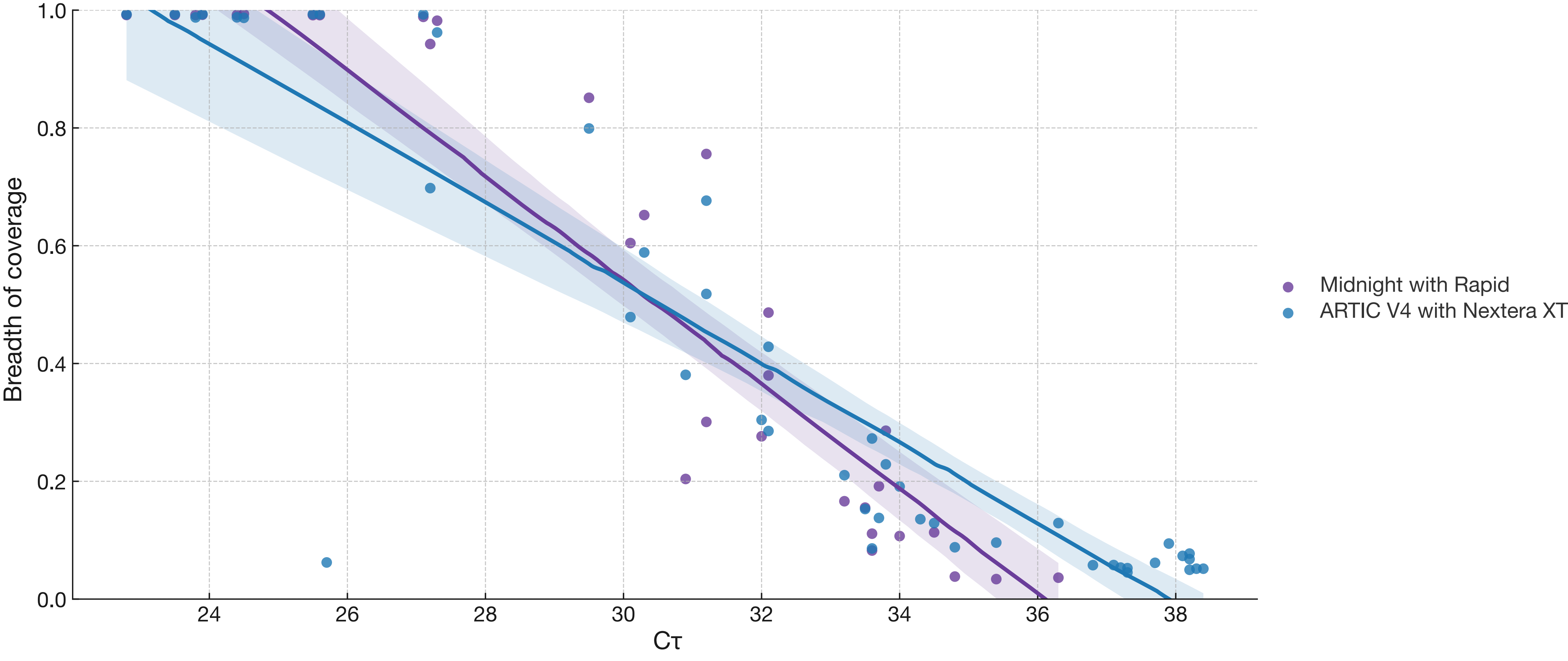
